# Flexible thin-film Implant with Depth Selectivity for Intraspinal Microstimulation

**DOI:** 10.64898/2026.02.23.707511

**Authors:** Soroush Mirkiani, Lukas Matter, Amin Arefadib, Katalin Sari, Neil Tyreman, Maria Asplund, Vivian Mushahwar

**Author notes:** Equal contribution to first authorship (Soroush Mirkiani, Lukas Matter). Equal contribution to senior authorship (Maria Asplund, Vivian Mushahwar). Corresponding author: Vivian Mushahwar.

## Abstract

Restoration of motor function after spinal cord injury remains a major challenge, as existing neuromodulation strategies such as epidural stimulation suffer from limited selectivity. Intraspinal microstimulation (ISMS) offers higher spatial precision but has been constrained by manually fabricated microwire arrays that lack reproducibility, depth control, and mechanical compatibility with neural tissue. Here, we present flex-ISMS, a thin-film, polyimide-based ISMS array integrating 42 stimulation sites distributed across 14 flexible arms. Acute *in vivo* implantation into the lumbosacral enlargement of domestic pigs demonstrated depth-specificity, site-selectivity and near normal recruitment of motor units resulting in graded contractions in muscles controlling the hip, knee, and ankle joints, with ranges of motion and isometric force generation approaching levels seen during natural locomotion (*e.g.,* 40° and 30 N for knee extension). Importantly, electrodes separated by 500 µm evoked distinct responses, underscoring submillimetre-scale selectivity. The high flexibility allows the device to conform to the spinal cord while displacing tissue by only 40×8 µm per arm. Histological analyses showed that the 125 µm diameter tungsten insertion aid of the flex-ISMS arms produced minimal acute damage, indistinguishable from that produced by conventional 50 µm diameter microwires. These acute outcomes establish the surgical feasibility and functional capability of flex-ISMS, and provide the foundation for forthcoming chronic studies in spinal-cord-injured models.

## Introduction

Spinal cord injury (SCI) is a devastating condition that can significantly reduce the quality of life of the affected person^1^. Disruption of spinal cord circuitries can lead to numerous effects including the loss of motor and sensory functions, spasticity, autonomic dysreflexia, cardiovascular diseases, diabetes, and mental health disorders^2,3^. Despite extensive research, there is no cure for SCI yet. One promising strategy to restore lower limb movement after SCI is electrical stimulation of the lumbosacral spinal cord below the level of injury^4–7^. In this context, epidural spinal cord stimulation (eSCS) has demonstrated promising outcomes, enabling standing and stepping in both preclinical models and humans^6,8–13^. However, the efficacy of eSCS is constrained by limited activation selectivity due to the shunting effect of the cerebrospinal fluid (CSF)^14–16^. To overcome this limitation, intraspinal microsimulation (ISMS) has been proposed as an alternative^17^. Currently, microwire-based ISMS implants (wire-ISMS) represents the predominant approach^18^; however, these implants face challenges, including labor-intensive manual fabrication, lack of depth selectivity, and mechanical mismatch with spinal cord tissue^19^.

ISMS is more invasive than eSCS because electrodes are implanted directly into the spinal cord with tips reaching the ventral horns of the lumbar/cervical enlarement to activate motor-related circuitry^20–36^. Submotor threshold, tonic eSCS combined with intensive, weight-supported locomotor training has demonstrated recovery of standing, overground stepping, and even volitional movements in motor-complete cases of SCI^9,10,37^. Recently, spatiotemporal and phasic eSCS delivered through personalized paddle leads with enhanced dorsal root selectivity enabled restoration of standing and overground stepping in participants with motor-complete SCI even without locomotor training^11^. However, the poor activation selectivity of eSCS, especially in caudal regions where overlapping spinal rootlets hinder precise targeting^11,38^, necessitates prolonged mapping sessions to optimize electrode configurations, which are further compromised by relative movements between the spinal cord and paddle lead^11,38,39^. In contrast, due to the activation of intraspinal networks and fibers in passage, ISMS can activate a variety of functional movements with higher selectivity and at lower stimulation amplitudes than eSCS^18,22,40–42^. Importantly, the conserved anatomical and functional organization of spinal motor networks across species enable precise targeting with the penetrating electrodes^20,43,43–46^. ISMS elicts long periods of fatigue-resistant, weight-bearing overground walking in cats^17,24^. In addition, ISMS can restore overground walking after chronic complete SCI in cats^21^.

Conventional ISMS arrays are typically constructed from microwires that are manually assembled into arrays. This results in substantial variability across implants due to the absence of automated fabrication processes^19^. These arrays are further constrained by a single stimulation site at the tip of each electrode, which limits precise targeting in the dorsoventral plane. If an electrode fails to evoke the desired motor response, it must be extracted and re-inserted. Another drawback is the mechanical mismatch at the implant-tissue interface, which can cause a long-term foreign body response (FBR)^47–50^. Strategies to mitigate FBR include minimizing the implant’s cross-sectional area, employing materials with a lower Young’s modulus^49,51,52^, eliminating tethering^52,53^, and applying biocompatible, anti-biofouling, or low-friction coatings^54–56^. Smaller and softer probes help reduce inertial forces at the tissue-implant interface, thereby minimizing micromotion-induced damage^49,57,58^. To achieve depth control, multi-site ISMS probes on flexible substrates have been fabricated and tested. However, these implants remain bulky (85µm cylindrical^28,59^ or 135µm by 120µm rectangular cross-sectional dimensions^29^) and require separate fabrication followed by bonding to fragile lead wires prone to failure.

To address the limitations of previous ISMS implants, we here present a proof-of-concept thin-film, flexible ISMS array (flex-ISMS) based on polyimide. Polyimide has been extensively evaluated in both brain and spinal cord implants, demonstrating excellent biocompatibility and mechanical compliance^60–63^. Its compatibility with photolithographic microfabrication enables precise patterning, reproducible manufacturing, and integration of multiple stimulation sites, thereby enhancing targeting and selectivity^64^. This study investigated two primary questions: whether a flexible thin-film ISMS array can be feasibly implanted in a large animal model, and whether such an array can reliably generate functional movements. To this end, we fabricated a 42-channel, flexible ISMS implant and successfully performed acute implantations in the lumbosacral spinal cord of three female domestic pigs. The evoked movement types, generated ranges of motions and joint forces demonstrated high selectivity enabled by depth-controlled stimulation. Voltage transients during *in vivo* stimulation indicated that, to maintain stimulation within safe limits during chronic use, further design refinements and electrode material adaptations are necessary. Importantly, the acute tissue damage associated with flex-ISMS implantation using tungsten insertion aids was comparable to that of single-site wire-ISMS. Collectively, these results highlight the strong potential of flex-ISMS for translation to chronic preclinical models and, ultimately, humans.

## Results

The first practical design of the flexible ISMS array (flex-ISMS) comprised 14 flexible arms (40 µm wide, 8 µm thick), seven arms per side of the spinal cord spaced 6 mm apart (Figure 1). Each arm had three electrode sites (each 75 × 15 µm) sputtered with an iridium oxide film (SIROF), and spaced 500 µm center-to-center to permit depth-controlled stimulation. A sub-set of implants with three arms per side of the spinal cord were also fabricated. Strain-relief consisted of short serpentine regions within each arm and along the longitudinal access of the implant, aimed at facilitating implant adaptation to movements. A polyimide extension cable connected the implant to a custom printed circuit board (PCB). Soak tests confirmed the stability of the polymethylmethacrylate (PDMS) encapsulated junction (Supplementary Figure S5). A detailed description of the design process and various implant iterations is provided in the supplementary material (Figures S1, S2).

**Figure 1.**
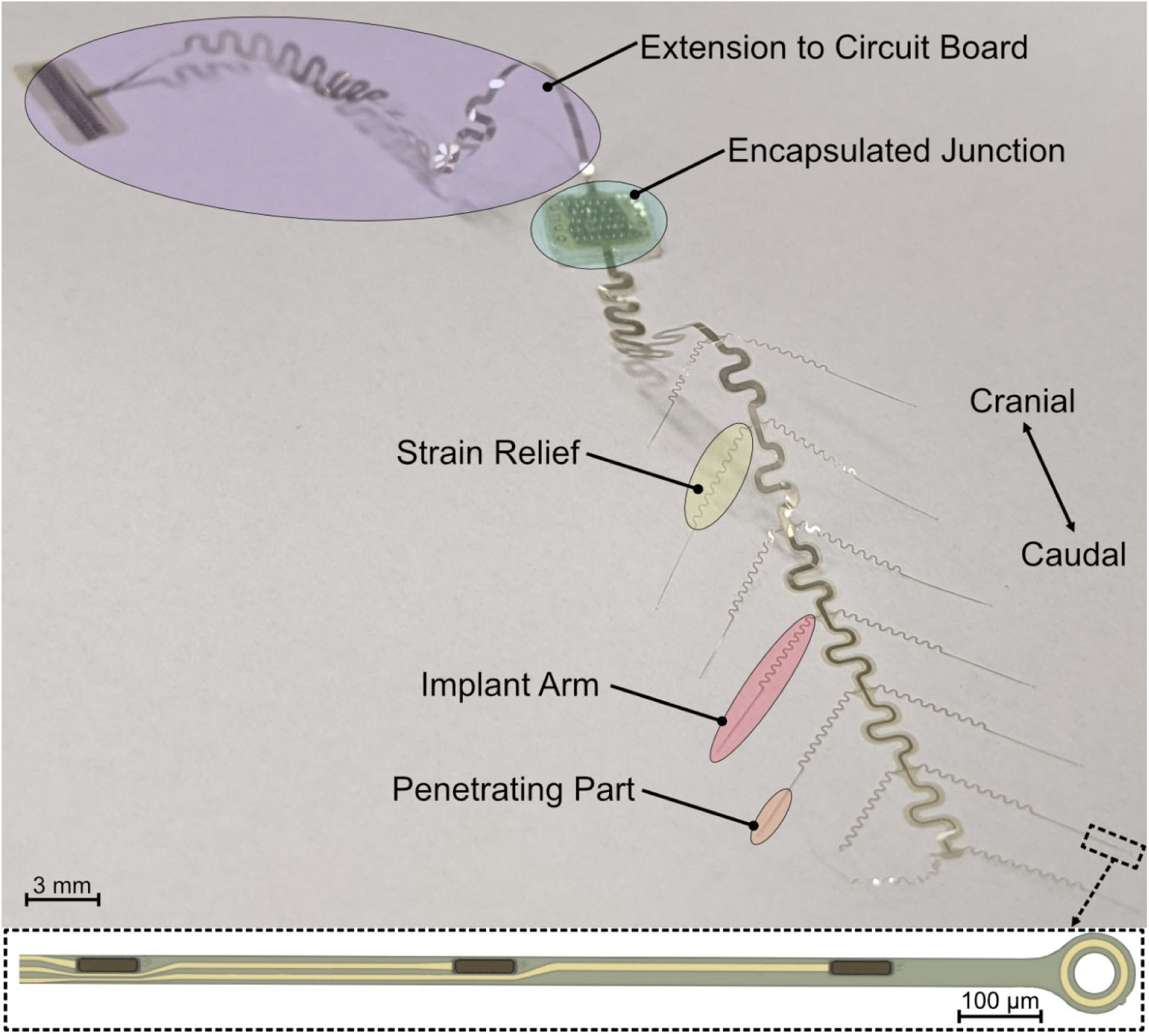
The flex-ISMS implant consisted of a connection point, serpentine sections to add strain relief and 14 arms with 6 mm separation. Each arm had 3 stimulation sites (70 x 15 µm) allowing for stimulation at 3 different depths (500 µm separation).

### Custom insertion aid for flex-ISMS implantation

Implantation of flex-ISMS required the development of a special insertion aid which we established through iterative trial and error (supplementary material, Figures S3, S4). Using a 125 μm tungsten rod with a 15° angle at the tapered 20 μm diameter tip as the insertion aid (Figure 2A), we successfully implanted flex-ISMS in three female domestic pigs (55.9 ± 1.0 kg) and demonstrated its ability to evoke discrete limb movements. For implantation, the laser-microfabricated insertion aids were threaded through the 50 µm diameter holes at the tips of each arm, then secured with polyethylenglycol (PEG; Figure 2A). The hole at the tip of each arm was designed with a ring of metal to mechanically reinforce the structure. The array was mounted onto a custom acrylic holder, designed to position the implant over the exposed spinal cord while preventing PEG dissolution upon contact with body liquids (Figure 2B). Individual arms were inserted sequentially with the tungsten aid advanced into the spinal cord and released upon PEG dissolution, leaving the implant arm in place. As shown in Figure 2C, flex-ISMS occupied negligible subdural volume and conformed closely to the curvature of the spinal cord surface. In total, 34 arms were implanted across three animals: six arms from the small array (3 per side) and 28 arms from two large arrays (7 per side each). All insertions successfully implanted the arm within the spinal cord, achieving a 100% implantation yield (each arm remained securely in place after withdrawal of the tungsten insertion aid). However, not all implanted arms elicited strong movements, likely due to suboptimal placement in the spinal cord.

**Figure 2.**
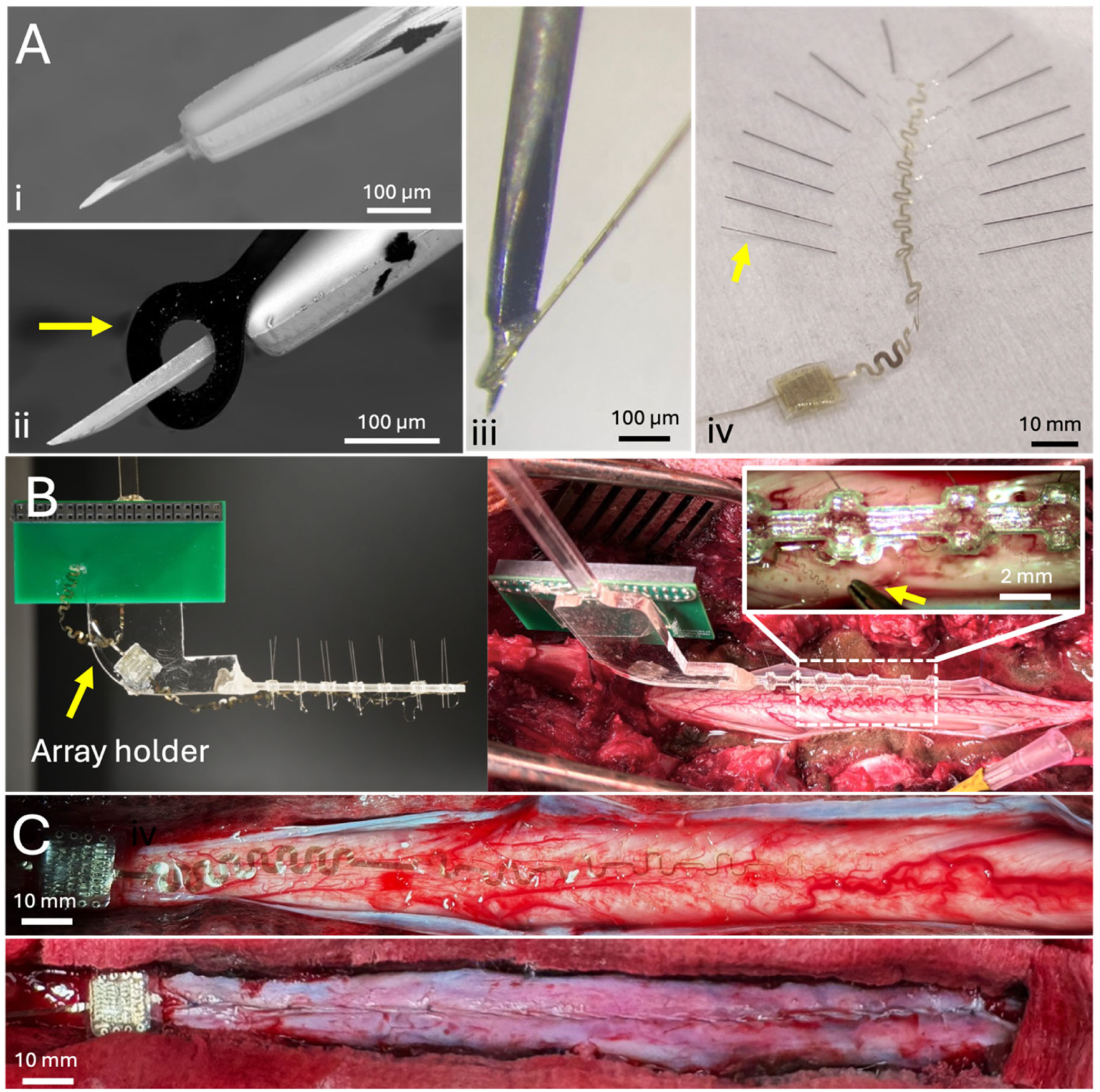
(A) Laser-micromachined tungsten insertion aids (yellow arrow) (i). Insertion aid tip is threaded through the insertion hole at the distal end of the flexible arms (yellow arrow in ii). Flexible arm is secured to the insertion aid with a small amount of polyethylene glycol (iii). All insertion aids are pre-attached to the implant arms to allow a fast insertion procedure (iv). (B) An array holder connected to a micromanipulator (left) is used for positioning the array on top of the lumbar spinal cord (right). Insertion aids are removed from the array holder and inserted one by one (right). The yellow arrow in the right inset shows an insertion aid detached from the array holder using a pair of forceps and ready for insertion. (C) The 8 µm thick flex-ISMS array conforms well to the surface of the spinal cord (top). Closure of the dura mater with the implant occupying the slightest space subdurally (bottom).

### Depth control in flex-ISMS enhances spatial selectivity of evoked movements

The flex-ISMS was evaluated using two array configurations: a smaller array with three arms, and a larger array with seven arms per side of the spinal cord. A total of 102 stimulation sites across three implants (34 arms) were tested. Detailed kinematic and isometric joint force analyses were performed on 30 sites (18 arms) that produced strong movements. Figure 3 shows representative types of evoked movements. The achieved movement types differed between array configurations and location of implantation along the lumbosacral enlargement. The small array, owing to its shorter rostrocaudal coverage, evoked knee and ankle extension and hip abduction (Figure 3A). The large array evoked a broader range of movements, including hip, knee, and ankle flexion and knee and ankle extension (Figure 3B, C). With the exception of a single bilateral hip abduction observed in one animal, all evoked movements were unilateral. This highlights the ability of the flex-ISMS array to target various locomotor-related regions within the lumbosacral enlargement to produce movements that constitute the gait cycle^65^.

**Figure 3.**
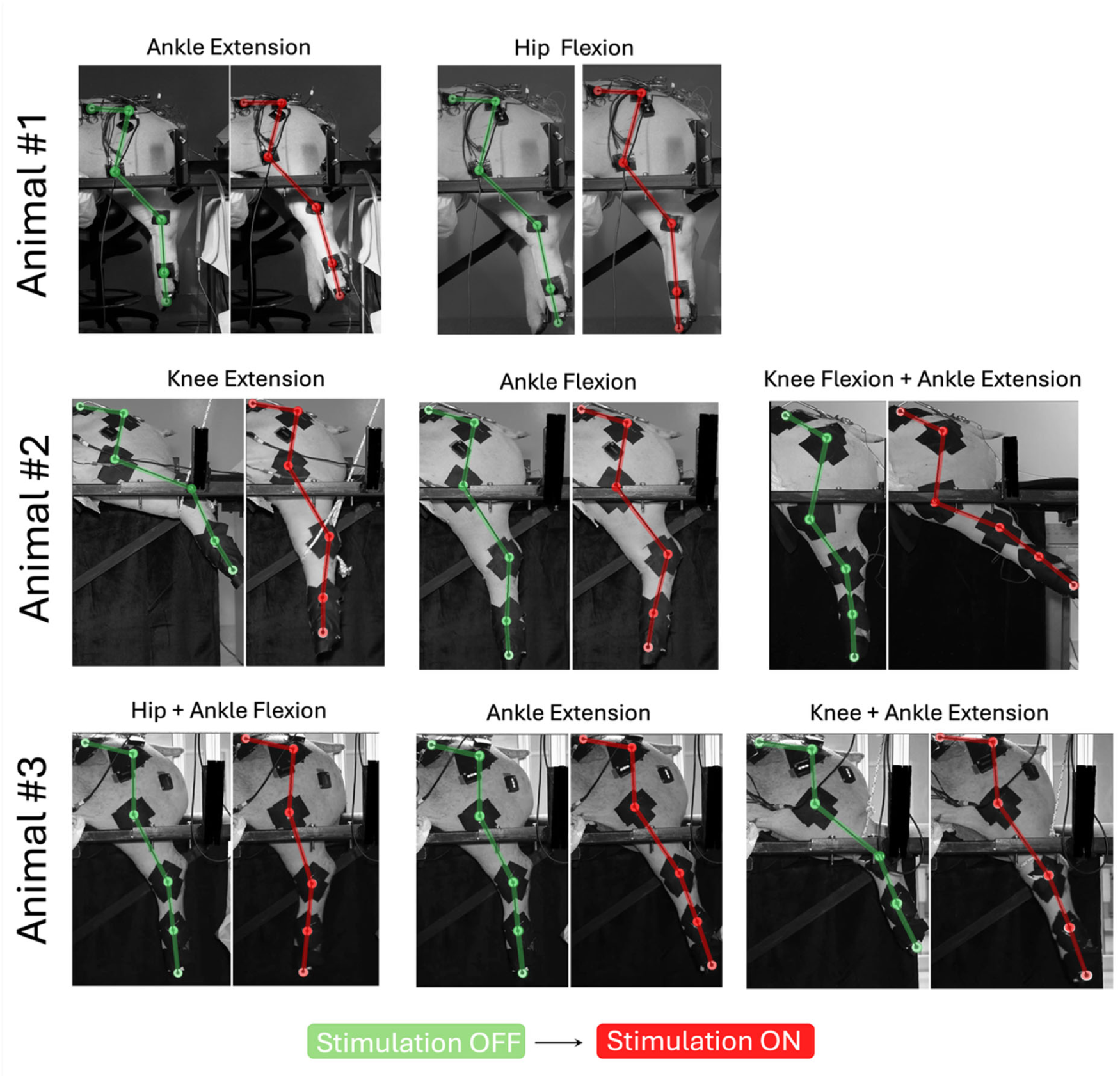
Flex-ISMS can generate various flexion/extension movements across the hip, knee, and ankle joints by targeting different locations inside the spinal cord of three animal. Animal #1 received the smaller array (3 arms per side) and could generate movements such as ankle extension, hip flexion, knee extension (not shown), and hip abduction (not shown). Animals #2 and #3 received the large array (7 arms per side). More diverse movements were achieved in these two animals as the array’s larger span could target more motor networks. Knee extension movements were recorded while the leg was pre-flexed against a counterweight. All movements shown were evoked by stimulation amplitude of 300 µA.

Figure 4 shows the range of motion (ROM) of the hip, knee, and ankle joints during movements evoked by stimulation through individual or multiple electrode sites on various arms of the larger flex-ISMS arrays, with stimulation amplitudes ranging from 25 to 300 µA. Stimulation through different electrodes on the same arm typically produced similar movements with varying strength. Nonetheless, distinct movement types (*e.g.*, a switch from flexion to extension across a joint) were occasionally observed when stimulating through different electrodes on the same arm (e.g., Figure 4A, ankle joint). To capture this site-specific response, joint ROMs were analyzed separately for each stimulation site rather than averaged across sites or animals. Stimulation depth-dependent effects were evident; in some cases, the movement strength increased as stimulation progressed from the dorsal to the ventral sites (Figure 4B), whereas in others the dorsal site produced stronger responses than the ventral site (Figure 4C, D, E). Simultaneous stimulation through multiple electrode sites on a single arm generally produced stronger movements. In some cases, combining electrodes resulted in an increase in the ROM of one joint while in other cases there was no change in the generated ROM. For example, in one case (Figure 4C), simultaneous stimulation of the dorsal and middle electrode sites increased knee flexion ROM from 4 ± 0.17° (dorsal site only) to 18.7 ± 1.1° at 300 µA. In another case (Figure 4B), stimulation through multiple sites yielded knee ROMs that were intermediate between the weaker (dorsal) and stronger sites (middle), while the ankle joint ROM remained closer to the stronger site.

**Figure 4.**
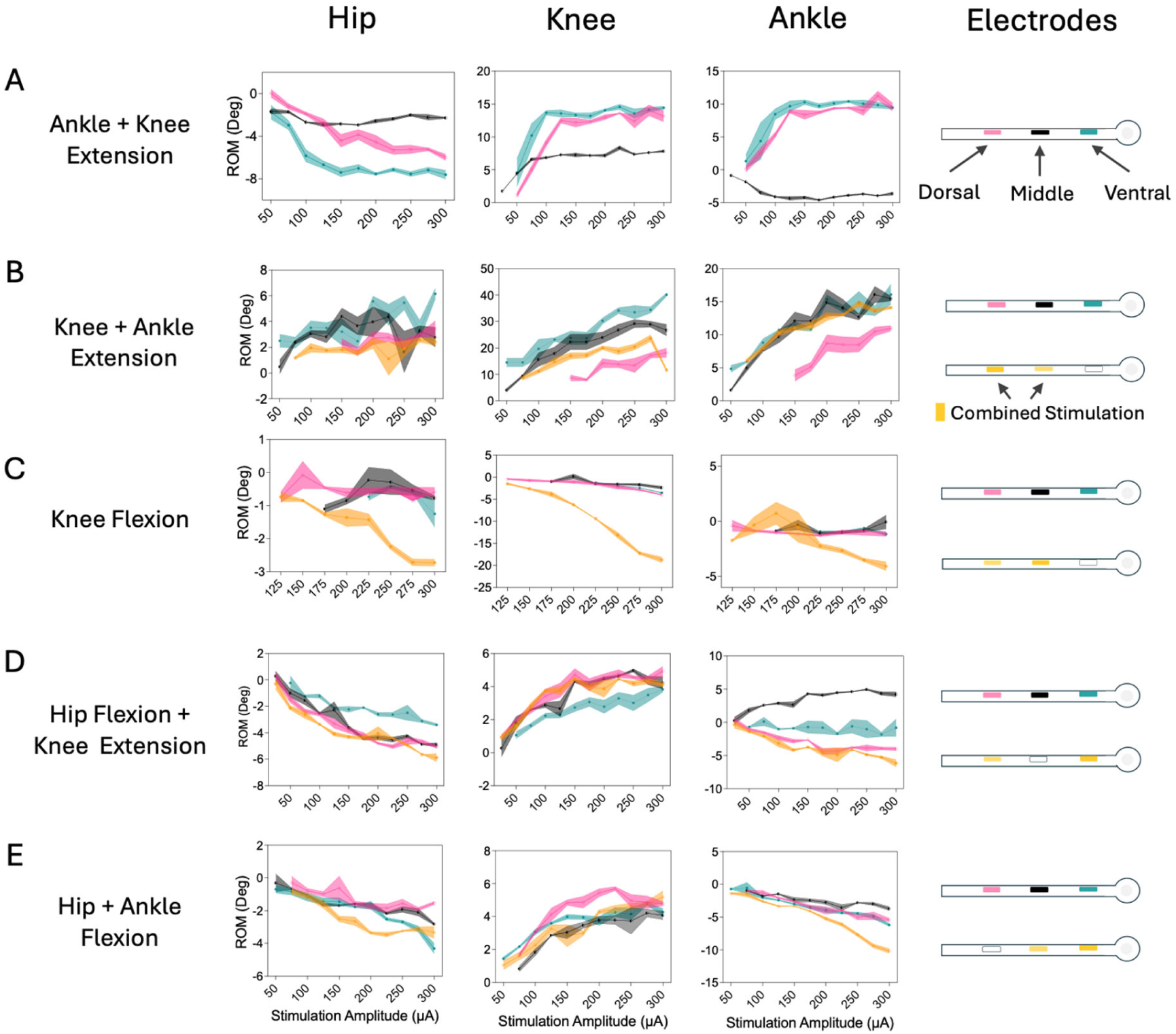
Examples of joint ranges of motion (ROMs) of induced movements by stimulation through single or a combination of sites simultaneously in two animals. ROMs of hip, knee, and ankle joints are plotted for stimulation amplitudes from 25 – 300 µA with three repetitions at each amplitude. Shaded areas show mean ± SE. Curves resulting from stimulation through individual or a combination of sites are color coded: ROMs resulting from stimulation through individual dorsal, middle, and ventral sites are plotted in pink, black and green, respectively. ROMs resulting from stimulation through a combination of sites are shown in yellow. Schematics on the right show the electrode sites for each evoked movement.

Figure 5 shows representative peak-to-peak isometric forces measured across a range of stimulation amplitudes (25 – 300 µA) for various stimulation sites. Force increased gradually with increases in stimulation amplitude at all stimulation sites (Figure 5). This graded recruitment of force is a key characteristic of ISMS^22,24^. Similar to the kinematics results, the magnitude of the generated forces varied across stimulation sties on each arm. In some cases, dorsal sites generated stronger forces (Figure 5B, C), whereas in others ventral sites generated stronger forces (Figure 5D). Knee extension forces of 19.7 ± 2.1N and 29.1 ± 1.8N were recorded from ventral and middle stimulation sites (Figure 5B) at 300µA, respectively. Combined stimulation of multiple electrodes resulted in higher (Figure 5E) or comparable forces to those generated by individual sites (Figure 5F). Shifting stimulation between sites on one arm altered both force magnitude (Figure 5B, C, E, F) and stimulation threshold for generating measurable force (Figure 5B,C,D). Moreover, electromyographic (EMG) recordings confirmed that site-specific stimulation could increase or decrease muscle activity, further supporting selective activation of distinct motor networks by flex-ISMS (Supplementary Figure S6).

**Figure 5.**
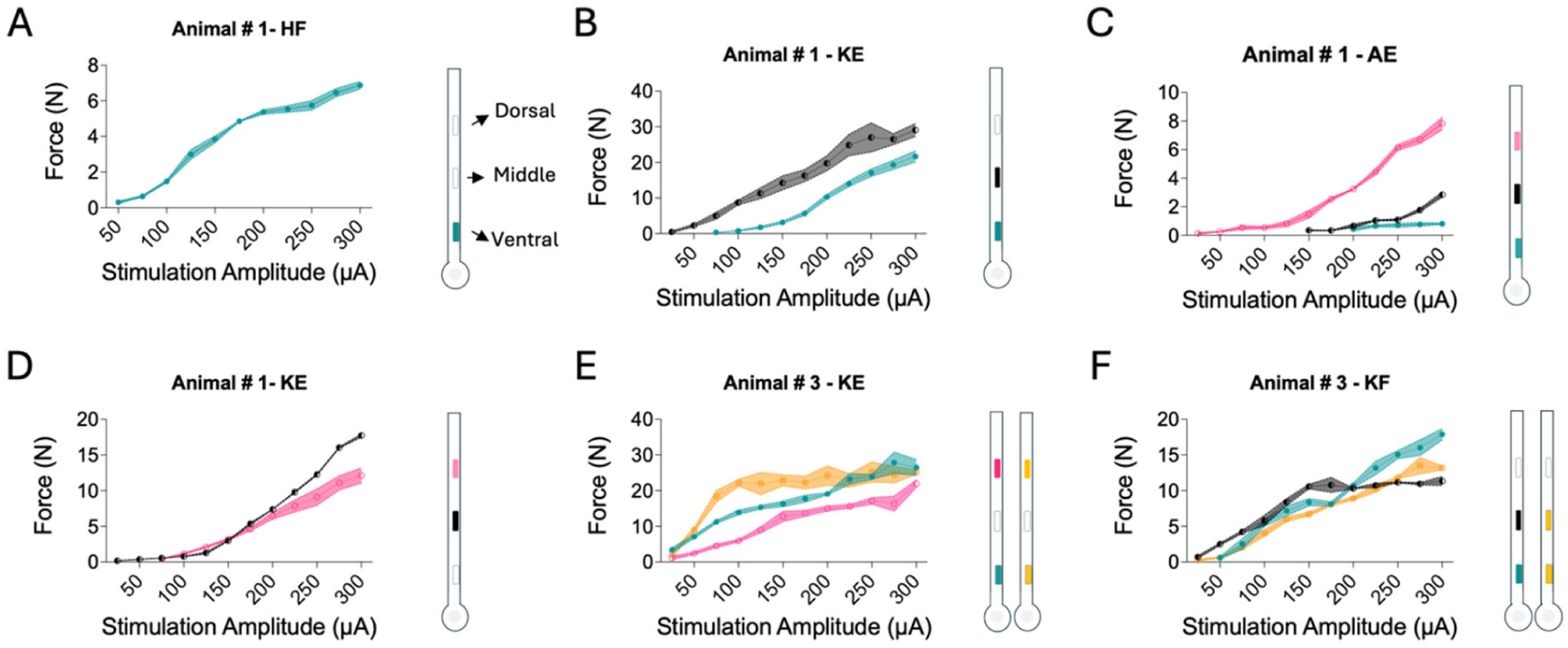
Examples of isometric joint forces recorded during evoked movements at stimulation amplitudes from 25 to 300 µA in three animals. The shaded area shows mean ± SD of force value at each amplitude with three repetitions at each amplitude. Isometric joint forces as result of stimulation through individual dorsal, middle, and ventral sites are shown in pink, black and green, respectively. Isometric joint force values from stimulation through combination of sites are shown in yellow. Schematics on the right side of each plot show the electrode sites stimulated and the configuration of combined stimulations. HF: hip flexion, KE: knee extension, AE: ankle extension, KF: knee flexion.

Together, the kinematic and force measurements demonstrate high spatial specificity of flex-ISMS. Specificially, the electrode spacing on each arm enables selective activation of locomotor-related networks, offering fine control over movement type and strength.

### Acute testing identifies charge injection requirements

The electrode sites of the flex-ISMS arrays were sputtered with SIROF. To support higher current densities, electrodes were coated with PEDOT/PSS^66^. The 1 kHz impedance magnitude of the PEDOT/PSS SIROF electrodes was 10 kOhm (Figure 6A) with charge storage capacity of 14 mC/cm^2^ (Figure 6C). Voltage transients (VT) were recorded to assess electrochemical stability and durability, yielding a safe charge injection limit (CIL) of 2.2 mC/cm^2^ at 125 µA (200 µs pulse width; maximum cathodic excursion E_mc_ < -0.6V) (Figure 6D). This is pulse-duration dependent, but largely in line with expectations based on what we previously have reported^67^. *In vivo* recordings at 25, 50 and 100 µA produced mean E_mc_ values of –0.48±0.2 V, - 0.56±0.1 V, and –0.93±0.2 V, respectively (Mean ± SD; Figure 6E,E and suplementary Figure S6). Since the *in vivo* VTs reflect the combination of voltage drop across tissue and electrode polarization, the actual electrode overpotential might be lower than the measured E_mc_. Under the tested conditions, ∼50 µA is thus taken as a conservative safe upper limit for current amplitude, assuming a symmetric biphasic pulse (Figure 6F).

**Figure 6.**
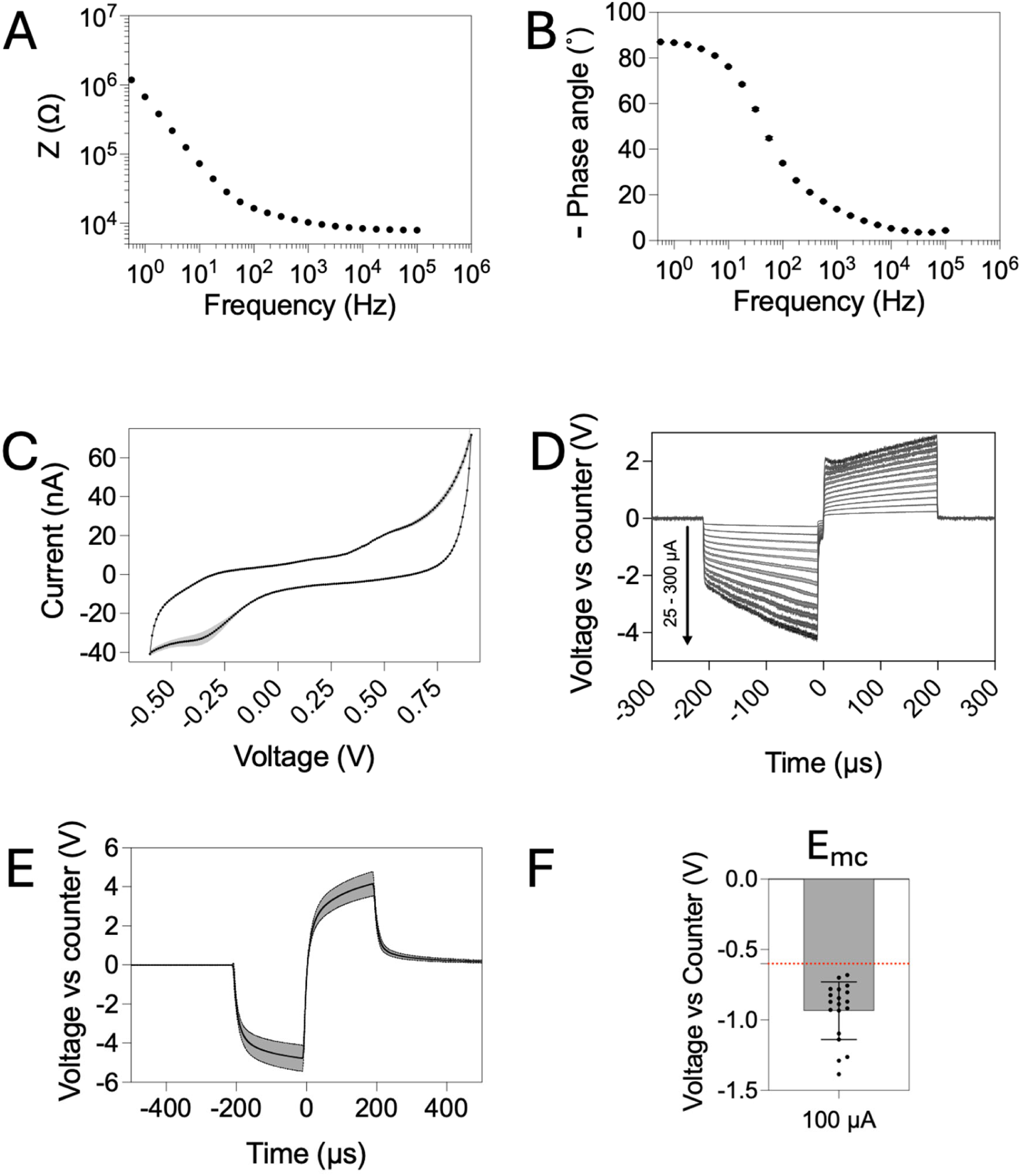
(A) Impedance, (B) phase angle, and (C) cyclic voltammetry at 100 mV/s sweep rate of exemplary PEDOT/PSS SIROF electrodes (n = 3). (D) Voltage transient experiments in 1xPBS aimed to investigate the durability of the electrode material showed that the PEDOT/PSS SIROF electrodes sustain a current of 300 µA (200 µs biphasic pulse) (n = 3). (E) In vivo voltage transient signals recorded in pig spinal cord at 100 µA from PEDOT/PSS SIROF electrodes. (F) the maximum cathodic excursion calculated from in vivo voltage transient responses at 100 µA; dashed line shows the –0.6 V safe limit (n=19).

The primary aim of this study was to explore functionality of flex-ISMS in acute experiments. Therefore, stimulation was extended up to 300 µA to probe different types of evoked movements, even though this amplitude exceeds the maximum safe cathodic voltage excursion. Nonetheless, this characterization provides an essential step that informs the required charge injection capabilities for eventual chronic use.

### Flex-ISMS implantation induces insertion damage similar to conventional wire-ISMS

Figure 7 shows representative hematoxylin and eoasin (H&E) stained sections after acute electrode implantation with either microwires (laser sharpened at 15°, Figure 7A, B) or flex-ISMS arms (Figure 7C, D) inserted using tungsten aids. The width of acute insertion damage was assessed in one animal using microwires and three animals using flex-ISMS. Although the limited sample size precluded statistical comparisons, the extent of damage was comparable for both wire- and flex-ISMS (Figure 7E). Averaged damage width measured in the coronal section showed higher values (∼200 µm) than those measured from axial sections (∼100 µm) for both implant types (Figure 6C). This discrepancy likely arises because in axial histological sections only a segment of the electrode tract is captured when it is not perfectly aligned with the cutting plane. Histological features were consistent with acute electrode insertion injury including the disruption of the neurovascular network, and increased blood–brain barrier permeability. These effects permit neutrophil infiltration and trigger upregulation of pro-inflammatory cytokines which in turn activate resident microglia^68–70^. Blood cells and microhemorrhages were visible in the electrode tracts of both wire and flex-ISMS. In some sections, multinucleated giant cells were visible digesting cell debris or red blood cells. The acute experiment time window (5-6 hours) was sufficient for early immune activation ^70^. To minimize tissue rupture during sectioning, implanted microwires were removed from the extracted tissue before freezing. In contrast, the thin-film flexible arms remained in situ and were sectioned together with the tissue, giving an impression of their dimension and placement within the insertion tract. Consequently, cross-sections of the sliced flexible probes could occasionally be observed in the coronal sections (yellow arrows in Figure 7D).

**Figure 7.**
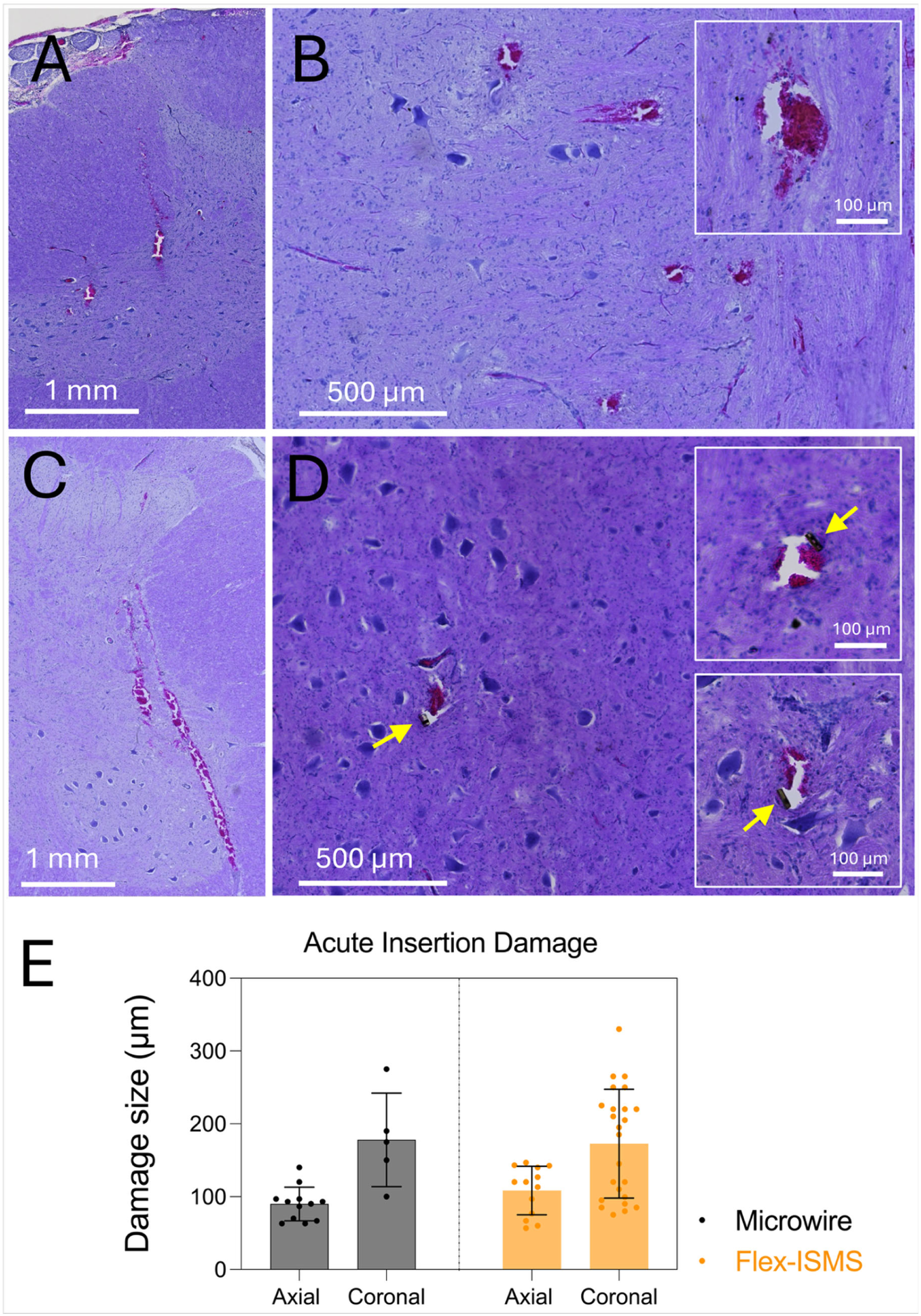
Examples of acute tissue damage from the insertion of 50 µm dia. microwire electrodes in axial (A) and coronal (B) sections stained with H&E. Example of acute tissue damage from implantation of flex-ISMS arm using 125 µm dia. tungsten insertion aid in axial (C) and coronal (D) planes. Localized hemorrhage together with disruption of neurons and axons along the insertion track can be observed in tissue. Insets show higher magnification images from coronal sections. Yellow arrows show cross sections of flex-ISMS arm cut with tissue. (E) Acute insertion damage width measured in axial and coronal sections (mean ± SD) from microwire implantation (one animal, n=12 microwire electrodes), or flex-ISMS arm implanted with insertion aid (3 animals, n=24 arms)

## Discussion

Flex-ISMS exhibits smaller foot-print, higher channel count, and greater flexibility than conventional microwire arrays. Existing ISMS implants have relied on single-site microwires (iridium or platinum-iridium, Pt-Ir)^19,21–23,30,71,72^, multisite silicon-based microelectrodes^43,73–76^, large-footprint flexible substrates^29^, or optical fibers^28^. Each approach presents limitations: single-site electrodes lack depth selectivity and require reinsertion if the desired movement is not achieved; multisite penetrating electrodes are either too stiff or too large and require additional lead wire connection points prone to failure^73,74,28,29^. In addition, previous multisite electrode arrays do not span the rostrocaudal extent required to target different locomotor-related neworks in the lumbosacral enlargement. Here, we demonstrated a proof of concept for flex-ISMS, a polyimide-based array featuring 42 electrodes distributed across 14 flexible penetrating arms seamlessly integrated to a a backbone structure aligned along the rostrocaudal axis of the spinal cord. The array is implanted bilaterally and depending on size, may provide both rostrocaudal and mediolateral coverage of the porcine lumbar spinal cord. Each arm holds three stimulation sites along its dorsoventral axis, allowing depth-selective stimulation. Through kinematic and isometric force analyses, we showed that flex-ISMS enables selective recruitment of joint movements and control over movement type and strength. In vivo VT measurements provided essential information for eventual chronic translation. Furthermore, a custom insertion strategy was developed, and histological analysis revealed acute tissue damage comparable to conventional wire-ISMS, despite the higher electrode count per implantation site. Importantly, flex-ISMS arrays could also be coupled with epidural cortical recording implants (used with existing eSCS platforms to predict motor intentions) to translate movement planing into modulation of flex-ISMS control program that targets the required networks with high specificity^77^.

Flex-ISMS enables joint movements with ranges comparable to natural locomotion. For example, knee extension of 40.1 ± 1.0° (Figure 4B) evoked from a ventral electrode site at 300 µA matched or surpassed joint excursions measured during slow overground walking in Yucatan minipigs (30–40° depending on speed)^65^. Another ventral electrode site produced a ROM of 19.6 ± 0.6° during ankle extension in a suspended leg. Adding this extension ROM value to the ankle flexion ROM obtained from stimulation through a ventral site (Figure 4E) of −6.2 ± 0.4° results in 25.8± 0.7° which is around the maximum ankle ROM during overground walking at 0.4-0.59 m/s (25.1 ± 3.1°) but lower than the ankle ROM when walking at 0.6-0.79 m/s (31.0 ± 2.5°)^65^.The limited hip extension observed may reflect that the flex-ISMS array did not extend caudally enough to target hip extensor muscle groups Nonethess, Hachmann et al.^42^ reported that the strongest hip extension movements are elicited from the L5 spinal cord segment, which was within the span covered by the large flex-ISMS array. Thus, inaccurate positioning of the flex-ISMS arms in this region may have resulted in the infrequent observation of large hip extensor ROM.

Isometric joint forces and joint ROMs increased nearly linearly with current amplitude, consistent with recruitment of motor units previously observed in rats^24^. The activation of axons of neurons in the vicinity of the electrode and fibers in passage can result in trans-synaptic activation of spatially separated motor pools a few centimeters away from the focal microstimulation^22,40^ resulting in motor synergies that are also activated during walking movements^20,35^.

Different intermediate and ventral regions in the spinal cord grey mater contain various neural populations^78^. Interneurons in the intermediate grey matter have direct connections to motoneurons that are involved in the generation of movements such as the flexor reflex^79^. More ventral regions contain interneuronal systems with abundance of fibers in passage projecting to motoneurons or other interneurons that are usually targeted by ISMS^17,20,40,80^. Activation of these ventral regions generates smooth movements often involving activation of groups of muscle synergies. Correct targeting of these overlapping intraspinal networks is crucial for successful clinical implication of ISMS. Given the dorsoventral extent of pig lumbar grey matter (≈4–5 mm) ^81^ and the ∼1 mm spacing between the most dorsal and ventral stimulation sites on the flexible intraspinal microstimulation (ISMS) array, the device can span intermediate to ventral neuronal populations.

PEDOT/PSS SIROF electrodes evoke strong movements despite their small geometric surface area (GSA). Each flex-ISMS electrode has a GSA of 1125 µm^2^, ∼40-fold smaller than conventional Pt-Ir microwires^22^. This is, in contrast to prior reports in pigs where activated iridium oxide film (AIROF) electrodes with conical tips and GSAs of 3000 µm^2^ were unable to generate movements unless stimulation was delivered through combinations of electrode groups simultaneously, and 24000 µm^2^ were needed to evoke movements^72^., In this study, stimulation through individual flex-ISMS electrodes was sufficient to produce strong movements. This discrepancy may reflect electrode geometry: planar electrodes provide uniform charge density distribution along the perimeter, whereas conical electrodes show higher charge densities at the tip and at the insulation-metal boundaries^82^. Comparable GSAs (1250 µm^2^ to 2400 µm^2^) have also been shown to produce hindlimb movements and bladder voiding^73,76^ in other species. The CIL of Pt-Ir microwire electrodes used for ISMS is reported as 60 µC/cm^2^ ^29^. Notably, SIROF coated with PEDOT/PSS demonstrated enhanced electrochemical performance, with an *in vitro* CIL of 2.2 mC/cm^2^. VTs recorded *in vivo* at increasing amplitudes indicated a lower limit of ∼0.8 mC/cm^2^. However, this reduction should be interpreted cautiously, as the *in vivo* VT recordings include the additional voltage drop across tissue, meaning the actual electrode overpotential is lower than measured. Such reduction of the CIL from *in vitro* to *in vivo* measurements have been reported previously and were hypothesized to be attributed to diffusional limitations in counterion availability^29,83,84^. It has also been suggested that electrode stability can be greater *in vivo*, owing to the adsorption of a protein layer that helps reduce dissolution, although such interfacial layers may also promote the onset of irreversible faradaic reactions^85,86^. Together, these findings highlight that multiple aspects influence electrode performance, making it difficult to define a universal method for determining the CIL. Prior experience suggests that the *in vitro* CIL can be used as an upper bound, with a safety margin applied when translating to *in vivo* stimulation^82^. Notably, flex-ISMS electrodes were at the edge of safe delivery of current amplitudes required by ISMS. Current amplitudes of around 100–150 µA are needed to evoke functional motor responses in pigs^23,29^. However, the CIL can be enhanced by changes to the design (*i.e.*, multilayers^62^) which would allow larger electrodes or a thicker PEDOT/PSS coating^87^.

The thin-film flex-ISMS design was motivated by the need to reduce the mechanical mismatch between the penetrating arms and the spinal cord tissue to reduce chronic FBR. Micro- and macro-scale motions can generate shear and compressive forces from the implanted shank that contribute to inflammation and glial scarring^47,50,57^ with severity influenced by several parameters, including cross-sectional area, stiffness, and tethering^49,51–53,88^. We replaced 50 µm diameter Pt-Ir microwires^23^ (1964 µm^2^ cross-section) with 40×8 µm polyimide arms (320 µm^2^ cross section). Even when the polyimide arm bends about its stiffer 40 µm axis (supplementary Figure S8B), it is ∼400-fold more compliant than the 50 µm dia. Pt-Ir microwires used in earlier ISMS arrays. This approximation is based on the fact that bending stiffness scales with EI (E: materials’ Young’s modulus, I = πr^4^/4 for cylinders; I = bh^3^/12 for rectangles) considering the contribution from the metal traces to be negligible. Bending about the 8 µm axis would make the arm more than 10,000-fold softer than the 50 µm dia Pt-Ir microwire. This order-of-magnitude reduction in flexural rigidity should markedly diminish chronic micromotion forces in the spinal cord. While this compliance prevents penetration through the porcine pia without support, insertion may be reliably achieved using sharpened 125 µm diameter tungsten rods. Smaller tungsten rods (75 µm or 50 µm diameter) did not provide enough bending stiffness at length of 25 mm making implantation challenging.

Although the 50 µm diameter Pt-Ir microwires are much smaller than the 125 µm diameter tungsten insertion aids, the measured size of the acute insertion damage was comparable for both cases (Figure 7C). A larger cohort is required to confirm this trend statistically, yet the tissue’s inherent elasticity likely allows the insertion track to rebound once the aid or electrode is withdrawn, minimizing residual damage. Prior studies indicate such acute lesions can resolve within weeks without persistent implant and interfacial stress ^89,90^, suggesting that thin-film arms with a smaller cross-section and high flexibility are likely to reduce chronic FBR. Nevertheless, long-term studies are required to confirm biocompatibility and durability.

## Future perspectives

This feasibility study has several constraints and we propose technology improvement in specific domains to enable chronic studies with implants. (i) Devices were patterned on 4-inch wafers, necessitating fabrication of a separate extension cord. Fabrication on larger diameter wafers would allow longer, fully monolithic arrays, and remove the need for the additional junction. (ii) Unlike recording probes, stimulation requires a large GSA for safe charge injection. This is limited by the present arm width and the number of stimulation sites that can be achieved. Multilayer metallization can be implemented in the future to introduce more stimulation sites and deliver the needed GSA while keeping the arms narrow. (iii) The integrated serpentine was not mechanically characterized in the current study; future work should combine finite-element modelling with bench testing to verify that it truly mitigates tether forces on the implanted arms. (iv) Manual implantation is operator-dependent, and the array holder used in this study does not allow real-time ultrasound guidance; a slimmer holder and robot-assisted insertion with live imaging in the future should improve the accuracy of implantation. (v) Lastly, we assessed only acute feasibility of the flex-ISMS arrays in this study. Long-term stability, electrochemical durability and tissue response remain to be established before thin-film flex-ISMS can be considered for chronic neuroprosthetic use.

## Conclusion

This work introduced flex-ISMS, a thin-film, polyimide-based, depth-selective ISMS array. Owing to its flexible polyimide substrate and PEDOT/PSS coated SIROF electrodes, flex-ISMS combined high conformability, large charge-injection capacity, and a 6-fold smaller penetrating cross-section than wire-ISMS while delivering more than twice the channel count and occupying minimal subdural space. Stimulation through individual sites elicited hip, knee, and ankle movements whose joint range of motion and isometric force scaled gradually with current amplitude. Electrode sites separated along the dorsoventral axis produced distinct movement patterns and strengths, highlighting the advantages of multiple stimulation sites. These results establish the acute feasibility of flex-ISMS. Future studies should focus on its long-term safety, stability, functionality, and therapeutic potential for translation to humans.

## Methods

All in vivo procedures were performed in accordance with protocols approved by the University of Alberta’s Animal Use and Care Committee.

### Implant Fabrication

The implants were fabricated in the cleanroom of the Chalmers University of Technology. The process began with spinning a layer of polyimide (PI) (U-Varnish-S, UBE Industries Ltd., Japan) onto a 4-inch silicon wafer at 4500 rpm to yield an ∼4 µm thick PI layer. The PI was then cured in a N_2_ atmosphere at 300 Torr, following the manufacturer’s specifications (120°C for 1 h, 150°C for 30 min, 200°C for 10 min, 250 °C for 10 min, 450°C for 10 min; rate 1°C/min). After curing, a bilayer photoresist was spun onto the PI (LOR 3A, Amolf, NL and S1805, micro resist technology GmbH, Germany). The resist was exposed using a laser writer (MLA 150, Heidelberg Instruments Mikrotechnik GmbH, Germany) and subsequently developed (MF-CD-26, micro resist technology GmbH, Germany). Next, the PI was treated with O_2_ plasma (1 min, 500 mTorr, 100W, 40 sccm O_2_, Plasma-Therm Batchtop VII, Plasma-Therm LLC, USA). Following this 100 nm of Pt was deposited (Lesker PVD225, Kurt J. Lesker Company GmbH, Germany). After liftoff and a second lithography step, ∼100 nm of Ir followed by ∼700 nm of IrOx was deposited (18 mTorr, 100 W DC, 26 sccm Ar, 14 sccm O_2_, Nordiko 2000, Nordiko Technical Services ltd, UK). Another liftoff process was performed. Before spinning the top PI layer at 4500 rpm, the wafer was treated with O₂ plasma to clean and activate the surface to promote improved adhesion of the subsequent layer. Following curing, a thick positive resist (AZ 10 XT 520 cP, MicroChemical GmbH, Germany) was used as an etch mask. The resist was exposed with the laser writer and developed (AZ 400K 1:3.5, MicroChemical GmbH, Germany). The outline of the implant was etched in a reactive ion etcher (60 mTorr, 200 W RIE, 400 W ICP, 2 sccm Ar, 50 sccm O₂; Oxford PlasmaPro 100, Oxford Instruments, UK). Initially, a mask aligner (MA6/BA6, Carl Zeiss AG, Germany) was used for exposure instead of a laser writer. The implants fabricated using the mask aligner were used for phantom implantations and preliminary surgeries. With the mask aligner, instead of using a bilayer resist, a negative resist (ma-N 1410, developed in ma-D 533/S, micro resist technology GmbH, Germany) was used for the lithography step prior to the metal depositions. All other fabrication steps were similar.

### Implant Assembly

The wafer was submerged in ultrapure water over night which reduced the adhesion of the PI to the carrier wafer. The implants were then carefully peeled off the carrier wafer using tweezers. Next, the meandered PI extension cord was soldered onto the implant (LFM-65W TM-HP, Almit Ltd., UK). Subsequently, the meandered PI extension cord was soldered to a PCB. The PCB served as an adapter, allowing to connect to the implant via a 2 mm pitch header. Finally, the junction between the PI extension cord and the implant was encapsulated with silicone (DowSil 734, Dow Corning, USA). The silicone was cured in air for 2 days.

### Electrochemical Characterization

All electrodes were characterized with electrical impedance spectroscopy (EIS) and cyclic voltammetry (CV) in 1xPBS (P3813, Sigma–Aldrich) before implantation using an autolab potentiostat (PGSTAT 302N, Metrohm Autolab B.V., Germany). The electrochemical setups consisted of the electrode on the implant as the working electrode, a stainless-steel counter electrode (∼ 20 cm^2^), and an Ag/AgCl reference electrode (Ag/AgCl, BASI, USA). Before characterization, all electrodes were shorted and activated with 100 CV-cycles (−0.6 to 0.9 V vs. Ag/AgCl, 100 mV/s). PEDOT/PSS was polymerized electrochemically on top of the sputtered iridium oxide film (SIROF). For the PEDOT/PSS deposition, electrodes were put in an aqueous solution containing 5 mg/mL sodium polystyrene sulfonate (Poly(sodium 4-styrenesulfonate), Sigma–Aldrich) and 0.01 M 3,4-ethylenedioxythiophene monomers (3,4-ethylenedioxythiophene 97%, Sigma–Aldrich). The voltage was set to 0.9 V vs. Ag/AgCl, and the charge was monitored and cut off once it reached 300 mC/cm^2^. Following the deposition, implants were rinsed with ultrapure water, CV-cycled, and characterized.

Voltage transients were recorded in 1xPBS in a two-electrode setup, with the electrode on the implant as the working electrode, and a stainless-steel counter electrode (∼ 20 cm^2^). The pulse (200 µs width, balanced biphasic with 10 µs interpulse interval, 50 Hz, 25 – 300 µA amplitude) was generated by a stimulator (PlexStim, Plexon Inc, USA). The current and voltage were recorded with an Oscilloscope (DSOX1204A, Keysight Technologies, USA). In vivo voltage transients were measured in the pig spinal cord using a two-electrode setup: the implanted microelectrode served as the working electrode and a hypodermic stainless-steel needle inserted in an adjacent muscle acted as the counter electrode. Stimulation pulses (biphasic 200 µs long pulses, 50 Hz repetition rate, at 25, 50, and 100 µA) were delivered via an STG 4008 stimulator (Multichannel Systems GmbH, Germany), and current and voltage waveforms were captured on an MDO3014 oscilloscope (Tektronix, USA). The charge injection limit was defined based on the current amplitude at which the electrode’s maximum electrochemical potential excursions (E_mc_) reached –0.6 V during the cathodic phase.

### Surgical Protocols

Four neurologically intact domestic female pigs (54.45 ± 3.1 kg) and three freshly euthanized pig cadavers (∼50 kg) were used. Two cadavers were used to assess feasibility and develop implantation methods, insertion aids, and array design refinement (Supplementary materials). We first implanted different versions of polyimide arrays (4, 8, and 12µm thicknesses) with different dimensions without a metallization layer to refine the thickness of the array, and the shape of the insertion hole at the tip of each arm in the array. The third cadaver was used for evaluation and refinement of the array insertion aid holder design and implantation workflow (supplementary material Figure S3). For the live animal experiments, one pig underwent implantation feasibility and testing the insertion aids and collecting in vivo electrochemical voltage transients; the remaining three pigs were implanted to evaluate functional performance, including kinematic analysis, isometric joint force measurements, and EMG responses.

Pigs were premedicated with intramuscular ketamine (20 mg/kg) and glycopyrrolate (0.01 mg/kg), then anesthetized via continuous intravenous infusion of propofol (40–145 µg/kg/min), remifentanil (0.03–0.14 µg/kg/min), lidocaine (1 mg/kg/hr), and dexmedetomidine (0.2 µg/kg/hr). A brief isoflurane bolus facilitated endotracheal intubation. Animals were positioned prone on a custom surgical table with hindlimbs suspended to maintain neutral spinal alignment. Vital signs and reflexes were monitored continuously to ensure a stable surgical plane of anesthesia.

A midline dorsal incision exposed the L3–L6 laminae. After applying 2 % lidocaine to the periosteum, complete laminectomies of L4–L5 and partial laminectomies of L3 and L6 were performed to reveal the lumbosacral enlargement. A dural incision was then made, and the dura mater was tented to expose the spinal cord.

### Insertion aid Fabrication

The buckling of the flexible arms during implantation prevents pia mater penetration; Therefore, we developed insertion aids for implantation in the pig spinal cord. Tungsten rods (125 µm diameter, 2.5 cm length) were micromachined on a femtosecond laser system (Carbide, Optec, Belgium). Each rod was mounted in an acrylic block and cut to produce a 300 µm–long rectangular taper with either a 20 µm or 40 µm cross-sectional width. To achieve a square profile, the taper was cut, rotated 90°, and cut again. Laser parameters were: 343nnm (third harmonic), mark speed 40 mm/s, pulse width 243 fs, power 75%, and 150 repetitions per cut. Final sharpening at a 15° bevel was employed in identical laser settings but with 50 repetitions only.

### Implantation Procedure

Prior to array implantation, insertion aids were evaluated using individual polyimide probes fabricated on the wafer (mimicking the array’s flexible arms but without meandering geometry) in agarose (0.6%) spinal cord surrogates and a freshly euthanized pig cadaver. Flexible array arms were bonded to the selected insertion aid via polyethylene glycol (PEG). For array implantation, each arm was fixed to an insertion aid, then the assembly was fixed to an acrylic holder with thin strips of double-sided tape. Under a surgical microscope, the holder was aligned over the lumbar dorsal root entry zones using published anatomical landmarks and our spinal cord mapping data in domestic pigs^91^. A pair of forceps released the tungsten aid from the holder, and the aid was advanced into the cord. If tape adhesion impeded release, a drop of isopropyl alcohol was applied. Post-insertion, the aid remained in the spinal cord for 10–15 s to allow for PEG dissolution;, the aid was then retracted, leaving the flexible arm in situ. In cases of premature arm detachment (during holder release or early PEG dissolution), the arm was manually positioned atop the pia in the desired location and inserted using an insertion aid under high magnification surgical microscope (Leica, OHS1, Germany).

One freshly euthanized pig cadaver and four live pigs were used to evaluate array implantation methods. In the first live pig, we evaluated implanting the large array by placing it on top of spinal cord and and using insertion aids with 40 µm taper. Persistant mechanical attachment of the insertion aid tip to the insertion hole occurred often; therefore, insertion aids with smaller tips ( 20 µm) were employed for all subsequent experiments. Then we used the freshly euthanized pig cadaver to test the feasibility of a custom acrylic holder affixed to a micromanipulator in conjunction with a small array. The next three live pigs were implanted using the 20 µm taper and acrylic holder, the second pig with the small array, and the third and forth pigs with the large array. In every case, we implanted all arms on one side of the spinal cord before proceeding contralaterally. The junction at the extension cable and array was routed outside of the dural sac at the rostral end of the dural incision, and the dura mater was then sutured closed.

### Stimulation protocol and data collection

A hypodermic stainless-steel needle inserted in the paraspinal muscle was used as a return electrode. Stimulation was delivered at current amplitudes from 25 µA to 300 µA in 25 µA increments (200µA biphasic pulse train, 50 Hz, 500 ms duration) via a microelectrode stimulator (STG 4008, Multichannel Systems GmbH, Germany) to evoke specific flexion and extension movements for kinematic, force, and electromyographic (EMG) assessments.

Reflective markers were attached with double-sided tape to the iliac crest, hip, knee, ankle, and metatarsophalangeal joints, and a camera recorded their motion in the sagittal plane. For knee-extension trials, the knee were initially flexed using a counterweight-and-pulley system mounted on the surgical table. Marker positions were identified using threshold-based detection in custom Python scripts, and joint angles and ranges of motion were calculated from three repetitions per movement and then averaged for each animal. Kinematic data were collected across stimulation amplitudes from 25 to 300 µA.

Surface EMG wireless sensors (Delsys, Trigno Avanti, USA) were attached to vastus lateralis, biceps femoris, and gluteus medius muscles to capture muscle activity during each movement and at each stimulation amplitude.

Isometric joint forces were captured using a 150 lb load cell (Interface Inc., Scottsdale, AZ, USA) with custom amplification and digitized at 1,000 samples/sec on a Tektronix MDO3014 oscilloscope. For each movement and at each stimulation level (25–300 µA), forces were recorded in triplicate, and the peak force for each trial was extracted for analysis.

### Terminal Procedures and Perfusion

At the end of each experiment, deep anesthesia was confirmed by the absence of corneal and withdrawal reflexes, and animals were euthanized via bolus injection of pentobarbital sodium (Euthanyl, 100 mg/kg) through a jugular catheter. Immediately following euthanasia, the vasculature was flushed with 4 L of physiological saline and then perfused with 8 L of 10% formaldehyde via the carotid artery, allowing outflow through the hepatic vein. The L2–S2 spinal cord and surrounding tissues were then removed en bloc and post-fixed in 4% paraformaldehyde at 4°C for 24–48 h, during which post-mortem MRI scans were acquired. The spinal cords were then divided into blocks and cryoprotected by immersion in 10 % sucrose in PBS for 72 h, then 30% sucrose in PBS at 4°C until they sank (an additional 48–72 h). Tissue blocks were then embedded in optimal cutting temperature compound, snap-frozen in isopentane chilled on dry ice, and stored at –80°C.

### MRI Imaging

After euthanasia, the spinal cord was carefully dissected free of bone and submerged in Fluorinert (3M) to suppress susceptibility artifacts. High-resolution imaging was then performed on a Bruker BioSpec 3 T preclinical scanner (Bruker, Germany) using both a multi-echo 3D T2–TurboRARE sequence and a T2*-weighted gradient-echo (mGE) protocol at 0.125 × 0.625 × 1 mm voxel dimensions, all within a volume transmit/receive coil. T2*mGE was employed to detect the subtle susceptibility changes around electrode tracts, enabling visualization of microinjuries such as tiny hemorrhages or tissue disruptions that conventional T2 imaging might miss.

### Histology and Imaging

Serial transverse and coronal sections (20 µm thick) were collected on a cryostat, mounting three to five sections per SuperFrost Plus slide. Slides were air-dried for 30 minutes at room temperature before returning to –80°C. Each batch of ten slides encompassed 30–50 serial sections. From each batch, one slide was stained with Harris hematoxylin and eosin (H&E) to confirm electrode tract locations and to assess any tissue disruption arising from array implantation. Sections were imaged on a Leica DM4000 microscope (Leica Microsystems, Wetzlar, Germany).

## Acknowledgments

We thank the staff of the Ray Rajotte Surgical Medical Research Institute at the University of Alberta for their assistance with the surgical procedures. We also thank Dr. Richard Fox for his insights on device implantation. This work was funded in part by the US Department of Defense, the Canadian Institutes of Health Research, the Canada Foundation for Innovation, the Brain Canada Foundation, and the University of Alberta Hospital Foundation. SM was supported by an Alberta Graduate Excellence Award, a Faculty of Medicine and Dentistry Dean’s Doctoral Student Award, and an Alberta Innovates Scholarship. VKM is a Canada Research Chair (Tier 1) in Functional Restoration. This work was performed in part at Myfab Chalmers. LM and MA were supported by the Chalmers Gender initiative for Excellence (GENIE). LM was also supported by a scholarship from Chalmersska forskningsfonden.

## Notes

### Competing Interest Statement

The authors have declared no competing interest.

