## Supplementary Material for "Flexible thin-film Implant with Depth Selectivity for Intraspinal Microstimulation"

To contribute to ongoing research in the exciting field of ISMS, we here present our design iteration of the flex-ISMS array. Our objective was to develop a thinner variant of a laser-microfabricated silicone-based multielectrode array. The primary design requirements included: three electrodes per arm, seven arms per side to ensure sufficient coverage of ~2/3 of lumbosacral spinal cord, and a maximum arm width of 40 µm and thickness of 8 µm to minimize the FBR.

A key challenge was achieving broad spatial coverage within the limited area of a 4-inch wafer, which we used due to the optimization of our polyimide (PI) fabrication process for this wafer size. Although a 6-inch wafer would provide more space, transitioning to it would require process optimization, which we may pursue in future iterations.

Version 1 of the implant (Figure S1) involved constructing it from three separate components: top, middle, and bottom. Each component was composed of a base and projecting arms of varying lengths. This modular approach allowed for a greater number of implants to be patterned per wafer. Each arm was connected to a support frame via breakable hangers, designed to detach when the arm was inserted into the spinal cord. These hangers were implemented to prevent entanglement of the arms during handling and implantation. To assemble the device, the array was connected to an extension cable aligned using a custom 3D-printed base plate featuring alignment cylinders to the corresponding holes.

This initial design was fabricated as a passive structure, without any metal layer, for prototyping purposes. However, several issues emerged during evaluation:

- Lack of visibility under the surgical microscope: The absence of a metal layer made it difficult to visually locate the insertion holes. Manually coloring the PI structures with a black sharpie helped to some extent. The insertion hole size was too small even with the surgical microscope at highest zoom level.
- Arm release mechanism: Detaching the arms from the hangers required excessive bending. This could potentially be mitigated by reducing the width of the part of the hanger connecting to the arm and by reducing the length of the arm.
- Limited implantation flexibility: As the arms detached only upon insertion into the tissue, the implantation site was fixed, dragging the entire structure with the arm being inserted.
- Lack of strain relief: The design lacked features to accommodate spinal cord elongation.
- Assembly process: Mounting the multi-component array onto the 3D-printed base plate was time-consuming and labor-intensive, limiting scalability.

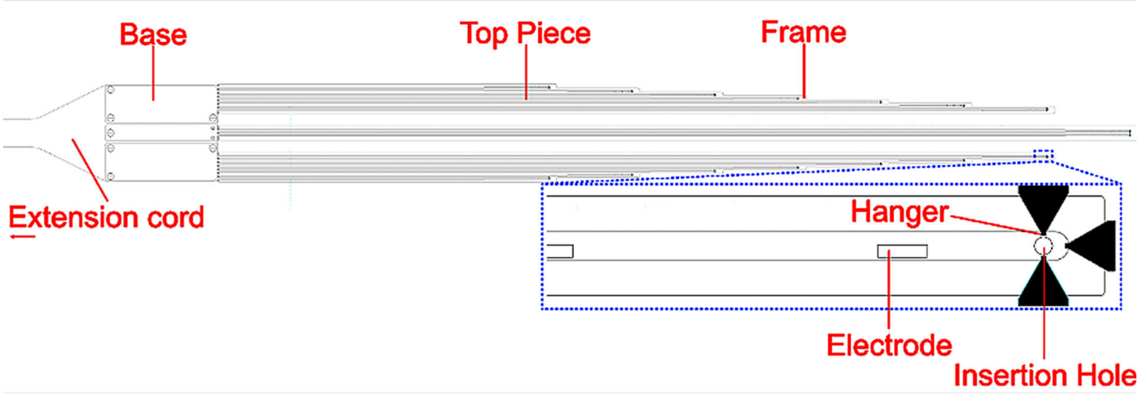


***Figure S1.*** *Version 1 of the flex-ISMS, featuring a modular structure consisting of three individual components with varying arm lengths, each connected to a central base via an extension cord. The arms are supported by a surrounding frame using breakable hangers, designed to detach upon insertion into the spinal cord through an insertion aid punching through the insertion hole.*

In version 2 of flex-ISMS (Figure S2), we introduced a central routing channel, in which all connection lines run through the middle of the implant. This configuration significantly reduced the required arm lengths compared to version 1, thereby facilitating arm release during implantation. While we retained the outer frame to prevent entanglement of the arms, we reduced the number of hangers to one per arm. Additionally, we incorporated meander structures in all parts. The base width was also reduced but it remained connected to an extension cord. The intended implantation strategy involved placing the implant, with its frame, directly onto the spinal cord. Insertion aids would then be used to sequentially release and insert the arms into the tissue. Perforations in the frame were designed to allow suturing the implant to the side walls of the opened dura mater .

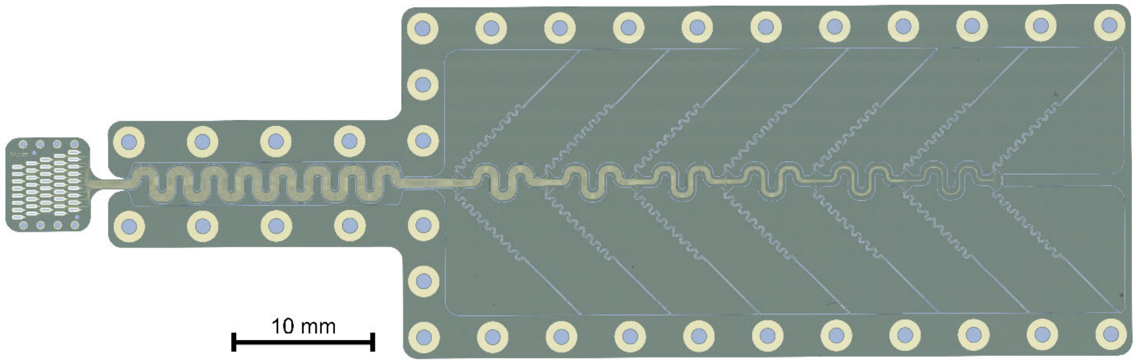


***Figure S2.*** *Version 2 of the flex-ISMS shown patterned on a 4-inch wafer. The design features a central routing of connection lines, shortened arms with integrated meanders, a reduced-width base, and an outer frame for spatial organization. One hanger per arm is used for temporary fixation, and suture holes are included in the frame.*

We evaluated version 2 in an agar plate to simulate implantation conditions. During testing, we observed that releasing the arms from the frame and subsequently inserting them into the agar remained challenging. Despite the reduced arm length and the use of only one hanger per arm, significant bending was still required to detach the arms, often causing them to cut into the agar. As a result, we decided to remove the external frame before implantation.

We tested the implantation in an acute pig experiment, however it was challenging to position the arms once they lay on to the cord. Once released, the arms tended to tangle and were difficult to visually identify and feed an insertion aid through the insertion hole.

To overcome these issues, we adopted a strategy in which the arms were preattached to the insertion aids using PEG. The insertion aids were then fixed onto a custom-designed acrylic holder, which could be positioned directly above the spinal cord. We first tested the smaller array using this technique in a freshly euthanized pig cadaver (Figure S3A). The initial tungsten insertion aid with a taper end that was 600 μm long and ~40 μm width was used in the cadaver experiment. This tapered end was too large and resulted in the insertion holes on the arm to remain mechanically attached to the arm post-insertion in the spinal cord (Figure. S4, right). The tip cross-section of the insertion aid was therefore reduced to 20 μm wide and ~300 μm long (Figure S4, left) for subsequent experiments. Furthermore, the design of the acrylic array holder was modified so that access to the insertion aids and their removal are improved. We used that approach for the remainder of the pigs included in the study.

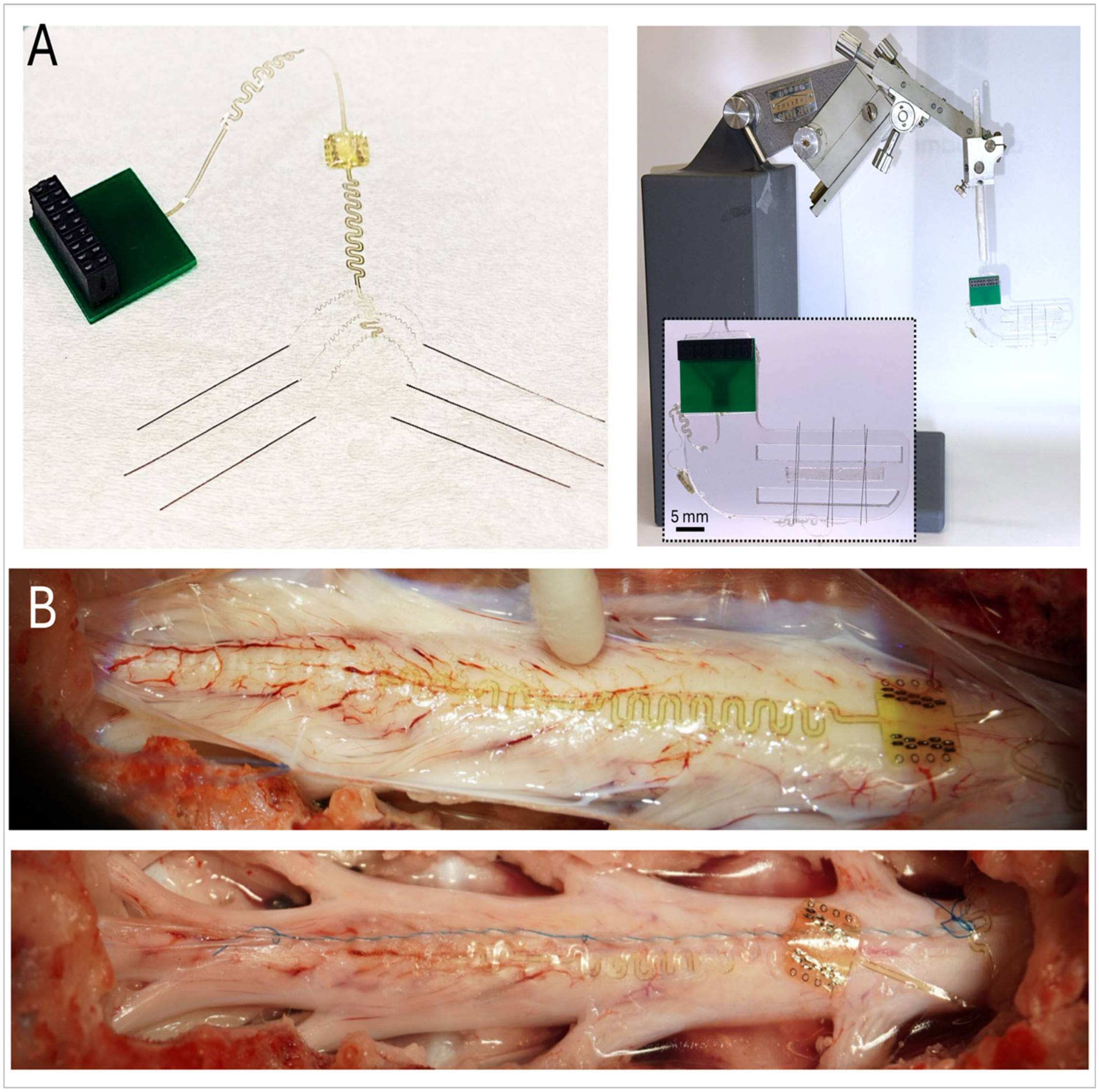


***Figure S3.*** *A) Arms of the smallr array are preattached to the insertion aids using PEG (left). Insertion aids are then attached to the acrylic array holder using double sided tape (right). The array holder is attached to a micromanipulator fixed to a base designed to deliver the array above the spinal cord during surgical implantation (B) The flex-ISMS array takes up negligible space in the subdural region (top) and the dura mater can be readily sutured back shut (bottom).*

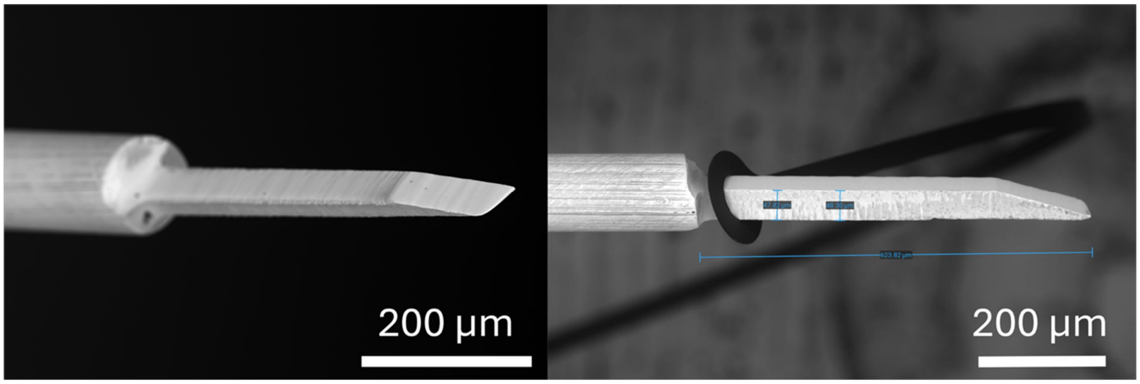


***Figure S4***. *Refined insertion aid (left) replaced the older design (right) that had little clearance between the tapered end as the holes at the tip of the array arms. often keeping the holes mechanically attached to the aid after implantation of the arm in the cord.*

*Encapsulation Test*

The aim of this study was to use flex-ISMS in acute experiments. We thus have chosen a simple encapsulation method for the connector junction by using DOW SIL 734. To evaluate the stability of the encapsulation prior to *in vivo* experiments, we first submerged the junction in 1xPBS at room temperature for 20 h. Electrochemical impedance spectroscopy (EIS) of electrodes was recorded hourly, showing no signs of encapsulation failure, with impedance magnitude remaining consistent throughout the test (Figure S5A). A side-view image of the encapsulated junction is presented in Figure S5B. To further assess resistance to water ingress, a second encapsulated device was immersed in 1xPBS at 37 °C for five days. EIS measurements were taken every 24 h on electrode E3, with the corresponding solder spot after the submersion experiment depicted in Figure S5C. The results confirmed the stability of the encapsulation, as the electrode remained connected throughout the testing period (Figure S5D).

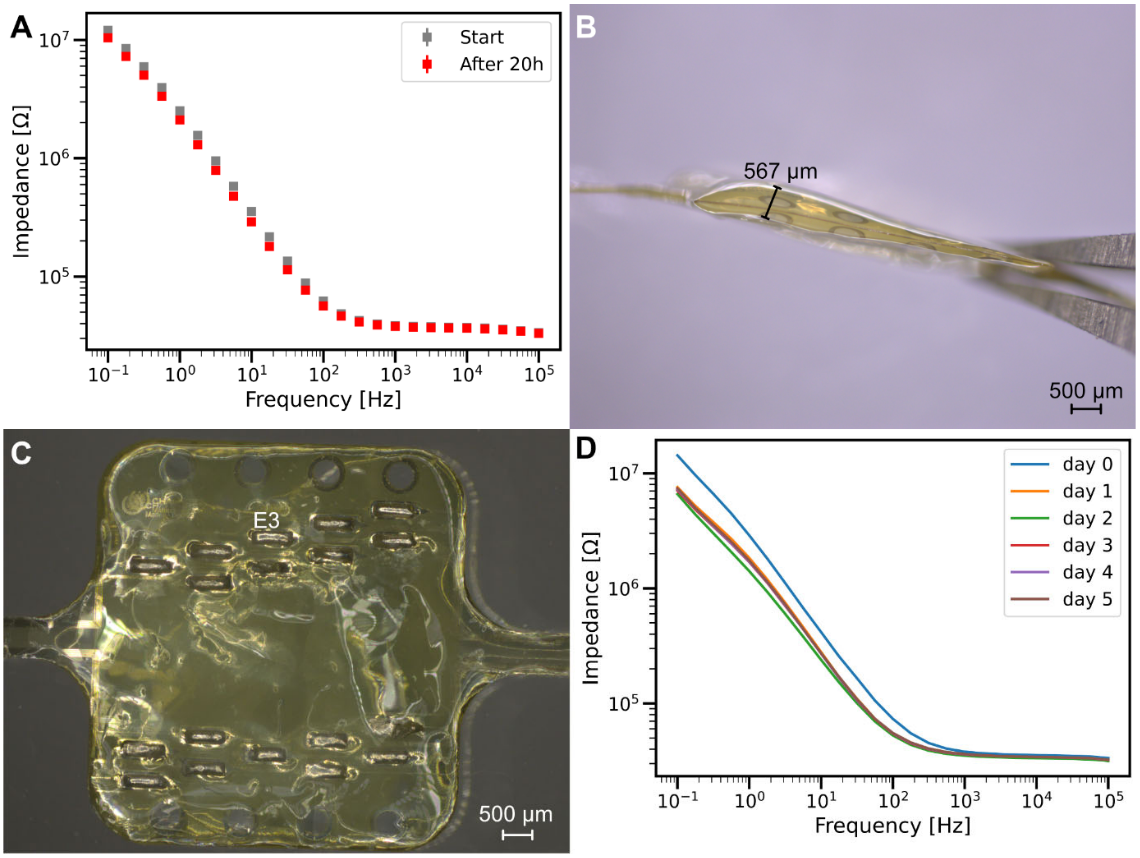


***Figure S5.*** *Evaluation of DOW SIL 734 encapsulation stability. (A) EIS measurements recorded hourly over 20 h with the encapsulated connector junction submerged in 1xPBS at room temperature showed a stable impedance magnitude, thus no signs of encapsulation failure (n = 3 electrodes on same implant). (B) Side-view of the encapsulated junction. (C) Image of the encapsulated junction after being submerged in 1xPBS for 5 days at 37 °C, with solder spot of electrode E3 marked. (D) EIS measurements taken every 24 hours over 5 days of E3 with the junction submerged in 1x PBS at 37 °C demonstrated a stable impedance magnitude, indicating effective encapsulation.*

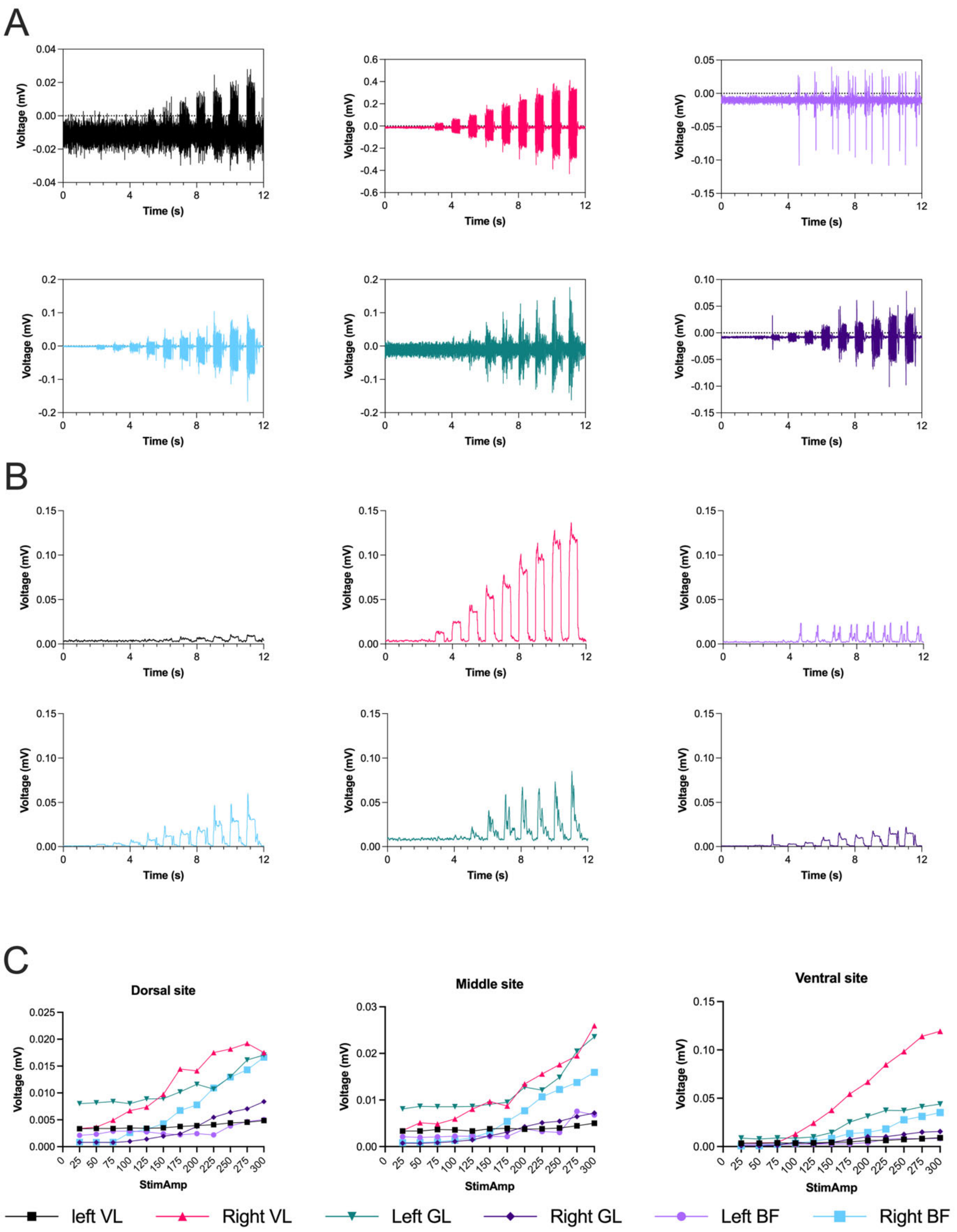


***Figure S6.*** *Representative electromyographic (EMG) activity recorded from 3 muscles bilaterally in response to stimulation through 3 electrode sites on one implanted arm at amplitudes ranges of 25 to 300 µA (0.5 s ON/ 0.5 s OFF) producing knee and ankle extension movement on the right side. (A) Raw EMG signlas recorded over the entire stimulation protocol from ventral site in the implanted arm. From left to right: vastus laterialis (VL), gluteus maximus (GL), and biceps femoris (BF). (B) Root mean square (RMS) envelope of filtered (20-450 Hz) signals. The RMS envelopes were computed using a 50 ms moving window applied to the squared filtered signal. (C) RMS amplitude calculated for each stimulation ON phase per amplitude level for 3 stimulation sites on one arm. EMG muscle activity amplitudes increase from dorsal to middle and to ventral sites.*

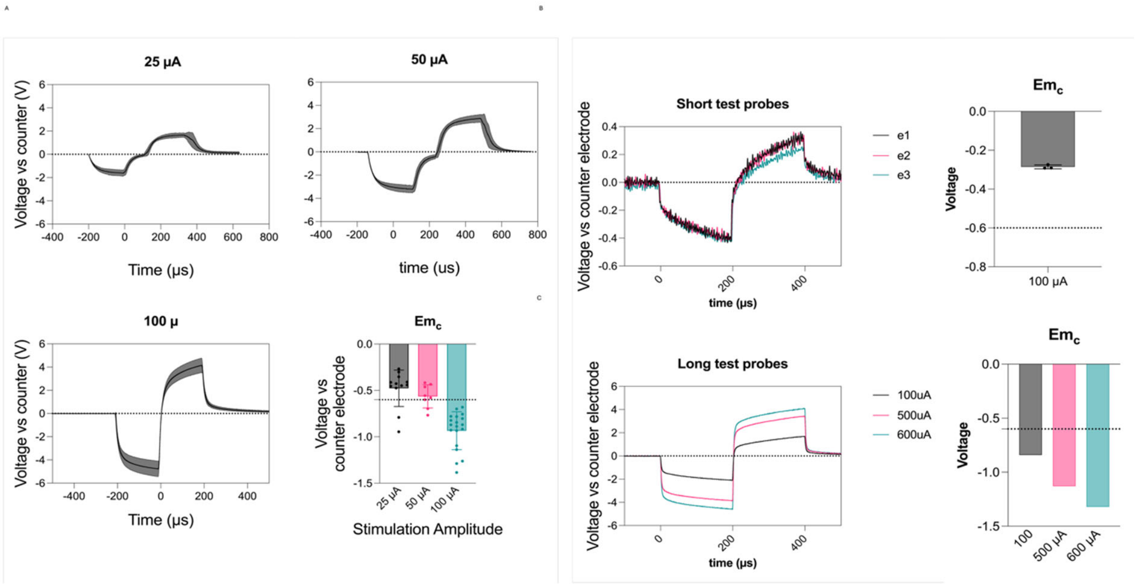


***Figure S7.*** *(A) In vivo voltage transient (VT) tests of flex-ISMS electrodes at 25, 50, and 100 µA amplitudes in 1 animal. Maximum cathodic voltage drop ($E_{mc}$) values are shown with respect to a stainless steel counter electrode. (B) Short test probes show smaller voltage drop than implant arm with longer tracks. (C) Longer test probes VT responses show higher access resistance because of longer connection lines and are stable during in vivo stimulation at amplitudes as high as 600 µA.*

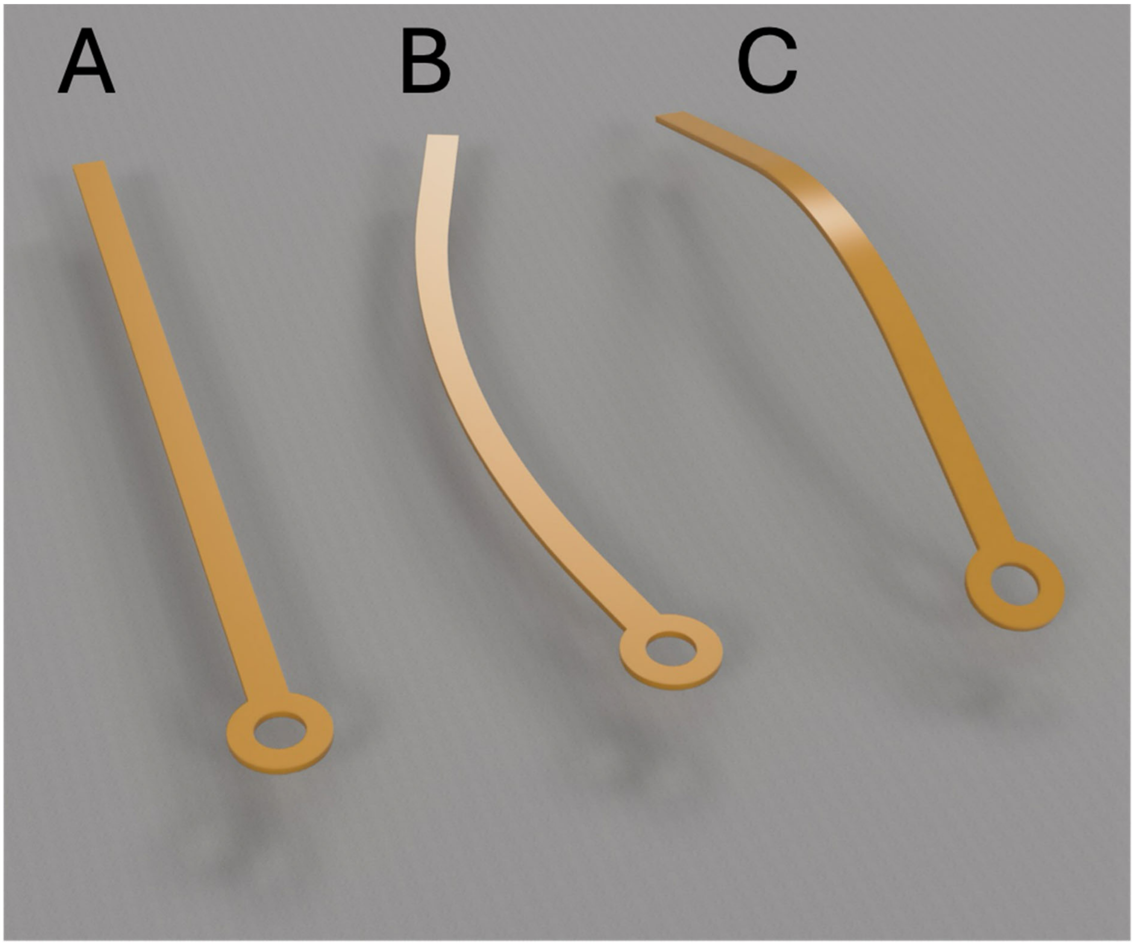


***Figure S8.*** *Schematic 3D representation of the flex-ISMS shank in its unbent state (A), when bent along its 40 µm width axis (B), and when bent along its 8 µm thick axis (C).*

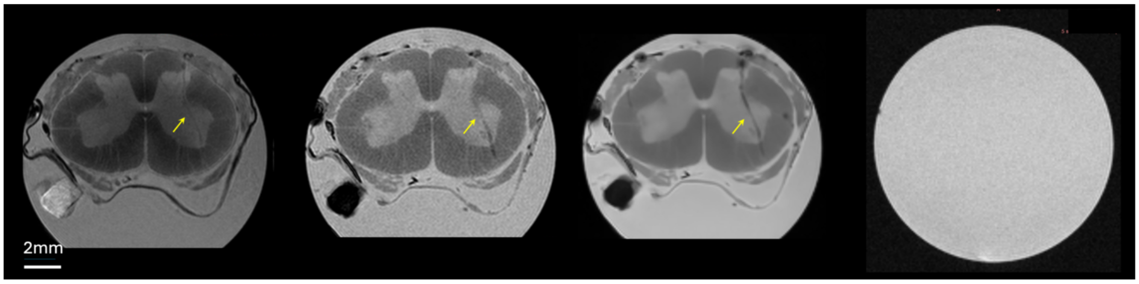


***Figure S9.*** *Example MRI snapshots from the stimulated spinal cord with flex-ISMS arms implanted. Three differernt sequences were tseted to enhance detection of slight susceptibility changes around electrode tracts (yellow arrows). From left to right; electrode tract using 3DT$_2$ Turbo RARE, FLASH (Fast Low Angle Shot), and T$_2$*-weighted gradient-echo (mGE). The far right snapshot shows an example slice of the flex-ISMS implanted in an agarose phanstom spinal cord showing no signal loss suggesting the artefacts seen in MRI are microhemorages in the electrode tract.*
